# Fast and accurate DNASeq Variant Calling workflow composed of LUSH toolkit

**DOI:** 10.1101/2023.03.01.530618

**Authors:** Taifu Wang, Youjin Zhang, Haoling Wang, Qiwen Zheng, Jiaobo Yang, Tiefeng Zhang, Geng Sun, Weicong Liu, Longhui Yin, Xinqiu He, Rui You, Chu Wang, Zhencheng Liu, Zhijian Liu, Jin’an Wang, Xiangqian Jin, Zengquan He

## Abstract

**Background:** Whole genome sequencing (WGS) is becoming increasingly prevalent for molecular diagnosis, staging and prognosis because of its declining costs and the ability to detect nearly all genes associated with a patient’s disease. The currently widely accepted variant calling pipeline, GATK, is limited in terms of its computational speed and efficiency, which cannot meet the growing analysis needs.

**Methods:** In this study, we propose a fast and accurate DNASeq variant calling workflow that is purely composed of tools from LUSH toolkit. The LUSH pipeline is highly optimized for the WGS pipeline based on SOAPnuke, BWA and GATK which can be deployed on any general-purpose CPU-based computing system. We validated the accuracy, speed and scalability of the LUSH pipeline on several standard WGS datasets.

**Results:** Our test results show that the LUSH pipeline and the GATK pipeline are highly consistent in terms of accuracy, achieving over 99% precision and recall on NA12878. For speed, the LUSH pipeline completes 30x WGS data in 1.6 hours, which is about 17x faster than the GATK pipeline. From BAM to VCF, LUSH_HC even takes only 12 minutes, about 76x faster than GATK. Moreover, the LUSH pipeline shows favorable scalability in terms of thread and sequencing depth.

**Conclusion:** The LUSH pipeline provides far superior computational speed to GATK while maintaining a high level of accuracy comparable to that of GATK, which greatly facilitates bedside analysis of acute patients, large-scale cohort data analysis, and variant calling in crop breeding programs.

## Introduction

With advances in sequencing technology and lower sequencing costs, whole-genome sequencing is playing an increasingly important role in single-gene disease screening or diagnosis, individualized cancer therapy, and pharmacogenomic screening [1–5]. Because it allows for the rapid, simultaneous detection of virtually all genes in a patient that may be associated with disease, which is particularly effective for patients with very rare or novel diseases, atypical clinical presentations, or prognostic responses [6–9]. However, the large volume of off-machine data from WGS presents new challenges in terms of analysis time and accuracy. Delayed clinical decisions may lead to severe morbidity or mortality, especially in acutely ill patients with potentially treatable genetic disorders [10, 11]. Therefore, rapid and efficient WGS analysis tools or pipelines are essential for timely molecular diagnosis, staging and prognosis, and pharmacogenomics-based guidance.

Currently, the GATK best practices proposed by Broad Institute are widely accepted standards for WGS variant calling pipeline [12]. It usually consists of several steps: preprocessing, alignment to genome, sort alignments, mark duplicates, base quality score recalibration, and variant calling, each corresponding to a recommended tool. However, this process requires tens of hours to perform analysis on a single set of WGS data [13], which cannot meet the current demand for urgent medical detection of patients with tumors or severe genetic diseases. To solve this problem, several ultrafast WGS analysis tools have been developed, such as Genalice [14] and Isaac [15], but the accuracy of these algorithms are not widely recognized or validated as they do not follow GATK best practices. Although sentieon DNASeq claims to follow the GATK algorithm, it requires a license to use it and needs to be built on a highly optimized backend [16], which restricts its widespread application. Recently, some WGS pipelines based on heterogeneous computing (such as GPU, FPGA, etc.) have been proposed. The representative one is DRAGEN [17], which uses an FPGA engine for the acceleration of computationally intensive operations. However, it usually requires extreme hardware support, resulting in limited generality and high cost. Therefore, there is still a need to develop more tools that are fast, accurate, and easy to access and deploy.

In this paper, we develop a novel, fast and accurate pipeline for DNASeq variant calling, consisting of multiple LUSH components. The LUSH pipeline reconstructs analysis tools SOAPnuke [18], BWA[19] and GATK[12] using C/C++, and employs a new parallel computing architecture. Several redundant intermediate steps of reading and writing the same file are eliminated in the LUSH pipeline, which greatly avoids unnecessary I/O throughput and improves CPU utilization. In the following, we validate the computational speed, accuracy and scalability of the LUSH pipeline on several datasets.

## Materials and Methods

### WGS benchmarking datasets

#### NA12878 (HG001) WGS data

Raw paired-end FASTQ files of NA12878 were downloaded from NIST’s Genome in a Bottle (GIAB) project at https://ftp-trace.ncbi.nlm.nih.gov/ReferenceSamples/giab/data/NA12878/. Then, 20X, 30X, 40X, 60X, 80X,100X data sets are obtained by down-sampling the original WGS data set under a series of gradient coverage. The gold standard truth variant calls and high confidence genomic intervals (NIST v3.3.2) were downloaded from https://ftp-trace.ncbi.nlm.nih.gov/ReferenceSamples/giab/release/NA12878_HG001/NISTv3.3.2/

#### “CHM-Synthetic-diploid” WGS data

CHM-Synthetic-diploid was constructed from the PacBio assemblies of two independent CHM cell lines using procedures largely orthogonal to the methodology used for short-read variant calling, which makes it more comprehensive and less biased in comparison to existing benchmark datasets[20]. Paired-end FASTQ files were downloaded from the European Nucleotide Archive with accession number ERR1341793 (https://www.ebi.ac.uk/ena/browser/view/ERR1341793). The benchmark truth call-sets and high-confidence regions of CHM-Synthetic-diploid were downloaded were included in the CHM-eval kit [20] and available at https://github.com/lh3/CHM-eval.

#### Two trios WGS data

This data set includes two son/father/mother trios of Ashkenazi Jewish (HG002/NA24385, HG003/NA24149, HG004/NA24143) and Han Chinese ancestry (HG005/NA24631, HG006/NA24694, HG007/NA24695) from the Personal Genome Project. Raw paired-end FASTQ files were downloaded from NIST GIAB repositories at https://ftp-trace.ncbi.nlm.nih.gov/ReferenceSamples/giab/data/AshkenazimTrio/ and https://ftp-trace.ncbi.nlm.nih.gov/ReferenceSamples/giab/data/ChineseTrio/. The truth call-sets and high-confidence regions used for benchmark were obtained from https://ftp-trace.ncbi.nlm.nih.gov/ReferenceSamples/giab/release/ with the latest version.

### Implementation of LUSH pipeline and GATK pipeline

The GATK pipeline was built according to best practices from https://gatk.broadinstitute.org/hc/en-us/sections/360007226651-Best-Practices-Workflows. Since most raw sequencing data require preprocessing operations to obtain clean data, such as removing adapters, low quality sequences and high N-base sequences, we include the SOAPnuke tool in the first step to preprocess the data, although this is not emphasized in GATK best practices. The LUSH pipeline followed a similar procedure as described for GATK best practices, including preprocessing, alignment to genome, sorting alignments, marking duplicates, base quality score recalibration and variant calling, consists of three LUSH components: LUSH_Aligner, LUSH_BQSR and LUSH_HC **(Figure 1, Table 1).**

**Figure 1.**
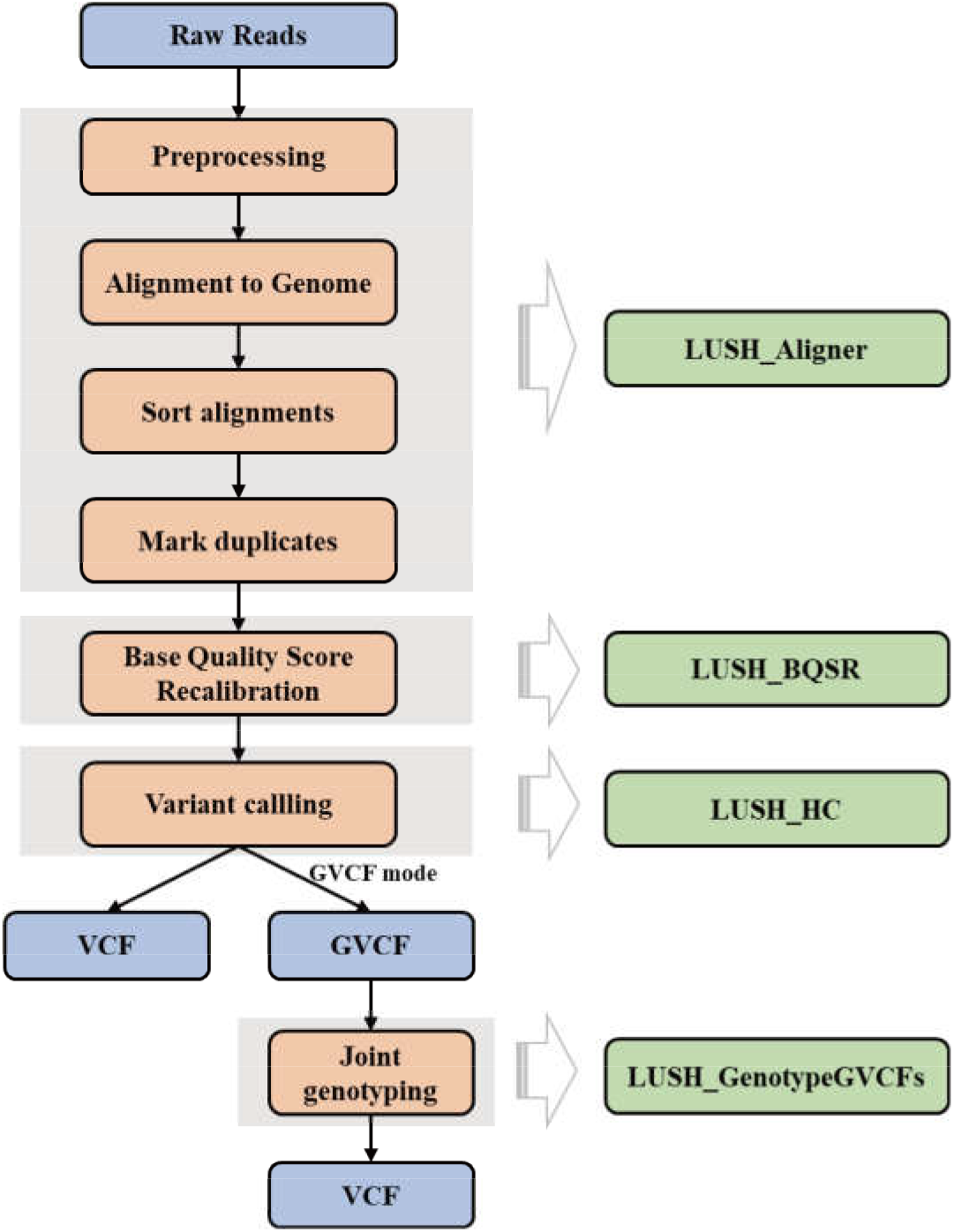
Overview of LUSH Variant Calling workflow. The LUSH DNASeq workflow is an optimized pipeline based on GATK best practices and consists of LUSH_Aligner, LUSH_BQSR, LUSH_HC, and LUSH_GenotypeGVCFs.

**Table 1.**
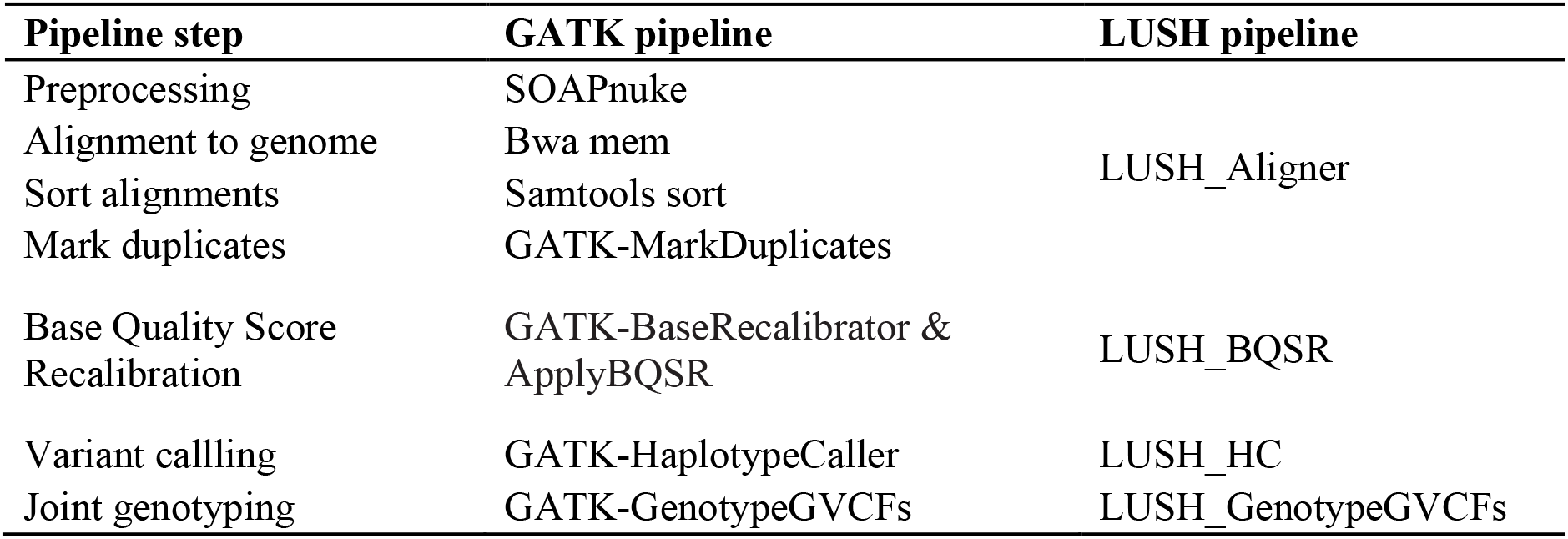
The composition of LUSH pipeline and GATK pipeline.

### Producer-Consumer pattern

The producer-consumer pattern is used in both LUSH_BQSR and LUSH_HC. With the buffered queue mechanism, each step runs independently, effectively improving the overall efficiency of the program.

The flow graph for LUSH_HC is shown in **Figure S1**. The main program starts by splitting BAM into distinct chromosomes, and each chromosome corresponds to a single producer thread. Within the given chromosome, the producer thread seeks for active regions that are possibly containing variants. Once one or more regions are found, they are pushed into the queue, waiting to be consumed by the consumer thread. Each consumer thread takes one region, and further calculation is done to call out variant genotypes in this region. Decoupling the detection of active regions from the main calculations for variant calling enables a manual control of number of threads at a time that are engaged in producers or consumers. A load balance is achieved to improve overall thread usage and hence accelerate the whole process.

### Computational Environment

All analyses were run on a Linux machine featuring an Intel(R) Xeon(R) Gold 6348 CPU 56-core processor and 500GB memory and were performed on a shared storage disk Dell EMC Isilon H500.

### Methodology evaluation

For NA12878 and Two trios WGS data sets, we used the haplotype comparison tool hap.py (v0.3.14, default comparison engine) for the comparison of diploid genotypes at the haplotype level to calculate the performance metrics. The variant calling accuracy of CHM-Synthetic-diploid WGS dataset was evaluated using RTG in CHM-evalkit (Version 20180222) [20]. The definitions of true positive (TP), false positive (FP), and false negative (FN) were based on the types of variant matching stringencies “genotype match”, and Precision, Recall, and F1-score were calculated as TP/(TP + FP), TP/(TP + FN) and 2*TP/(2*TP + FN + FP), respectively. Tools-specific SNPs and INDELs were annotated using the SNPEFF (v4.3) [21] with default parameters.

We evaluated the computational performance of all the tools on the same Linux machine, measuring both total runtime and maximum memory consumption using the “/usr/bin/time” command.

## Results

### Overview of LUSH DNAseq workflow

The LUSH DNASeq workflow is an optimized pipeline based on GATK best practices. Its main components are LUSH_Aligner, LUSH_BQSR, LUSH_HC, and LUSH_GenotypeGVCFs (**Figure 1**).

LUSH_Aligner integrates four functional modules: preprocessing, alignment to genome, sort alignments and mark duplicates, corresponding to the original software: SOAPnuke, Bwa mem, Samtools sort [22] and GATK-MarkDuplicates (Picard), respectively. LUSH_Aligner reimplemented the original software algorithm using C and C++ and optimized the algorithm architecture to improve CPU utilization while reduce IO usage. In the original WGS workflow, FASTQ data are preprocessed by SOAPnuke, including the removal of adapters, low quality sequences and high N-base sequences, and the final output of clean FASTQ files. Then, the FASTQ files have to be read again for genome alignment, and the following steps of sorting and marking duplicates also have the process of constantly writing and reading the same file, which greatly increases the disk I/O and slows down the workflow. The integration of LUSH_Aligner eliminates the repetitive process of writing and reading files. In the parallel architecture, the aligned results are split by genomic position and then sorted alignments and marked duplicates in multiple threads.

LUSH_BQSR is a C/C++ re-implementation of the GATK BaseRecalibrator and ApplyBQSR algorithms. It uses a parallel computing architecture based on the producer-consumer pattern, where the producer thread is responsible for data distribution and the consumer thread is responsible for data processing and computation (**see methods**). This pattern can effectively improve system performance as producers and consumers can run concurrently, thus increasing the efficiency of data processing.

LUSH_HC is a C/C++ re-implementation of the GATK HaplotypeCaller algorithms, which also uses a parallel computing architecture based on the producer-consumer pattern **(Figure S1, more detail see methods)**. LUSH_HC also implements the GVCF mode algorithm to meet the demand of GVCF files in cohort studies. Correspondingly, the C/C++ reimplementation of LUSH_GenotypeGVCFs is used to perform joint genotyping on one or more samples.

### Computational Performance on different threads and sequencing depth

The computational performance of the pipelines may not necessarily improve with the increase of the number of cores used. Application performance can be limited due to multiple bottlenecks including contention for shared resources such as caches and memory. Thus, we specified 12, 24, 36, 48 and 56 (max) threads at a single node to test the single-node scalability of the LUSH pipeline. As shown in **Figure2**, the runtime of all LUSH tools in this pipeline decreases significantly as the threads are increased. The pipeline completed in ~4.89 hours when running at 12 threads and ~1.6 hours when running at 56 threads (**Figure2 A**). Indicating that the LUSH pipeline has great thread scalability.

**Figure 2.**
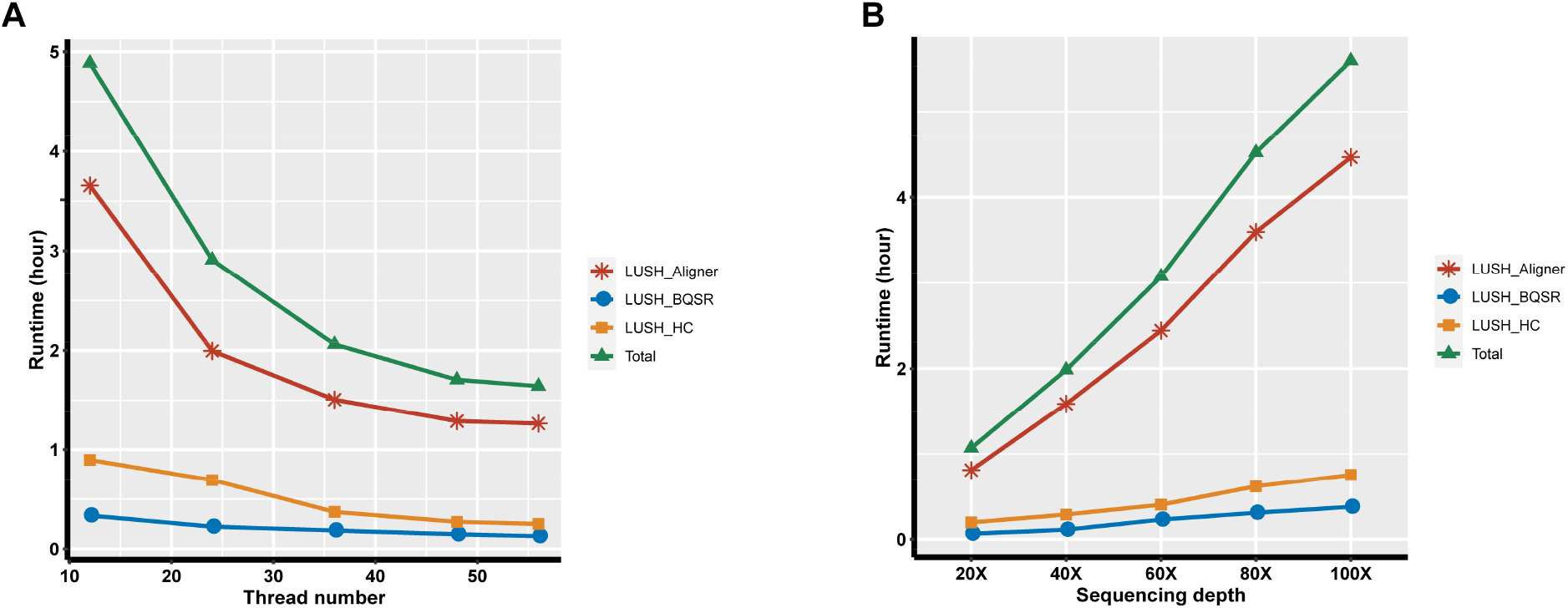
Computational performance of the LUSH pipeline at different threads and sequencing depth on NA12878. **A)** Running time with 12, 24, 36, 48 and 56 threads on NA12878 (30X). **B)** Running time at sequencing depth 20X, 30X, 40X, 60X, 80X,100X. Each data point is the average of two replicates.

We then generated NA12878 WGS datasets with 20X, 40X, 60X, 80X and 100X coverage depths by down-sampling. Each dataset was performed with the LUSH DNASeq pipeline to investigate the effect of sequencing depth on performance. Each task was run on the maximum available cores (56). The runtime of all LUSH tools and the entire pipeline increases almost linearly with increasing sequencing depth (**Figure2 B**).

### Speed of LUSH pipeline relative to GATK pipeline

To compare the performance of each component of the LUSH pipeline with that of the GATK pipeline, we analyzed the NA12878 30X sample on a maximum 56-core machine. Each software thread parameter was adjusted to the maximum available. LUSH_Aligner integrates four functional modules for the FASTQ to BAM process, including pre-processing, alignment to the genome, sorting alignments, and marking duplicates. LUSH_Aligner completed FASTQ to BAM on NA12878 WGS 30X in less than 1.3 hours, which is more than 5 times faster than the 6.88 hours of GATK pipeline (**Figure 3 A**). LUSH_BQSR integrates BaseRecalibrator and ApplyBQSR of GATK pipeline to greatly improve thread utilization. On the NA12878 WGS 30X, LUSH_BQSR took only about 5 minutes, which is about 60 times faster than the 5.22 hours of the GATK pipeline (**Figure 3 B**). To produce a VCF from a BAM file, HaplotypeCaller was widely recognized as the most time-consuming step in the GATK best-practice pipeline. It took ~15 hours to complete the NA12878 WGS 30X, while LUSH_HC took only about 12 minutes (**Figure 3 C**).

**Figure 3.**
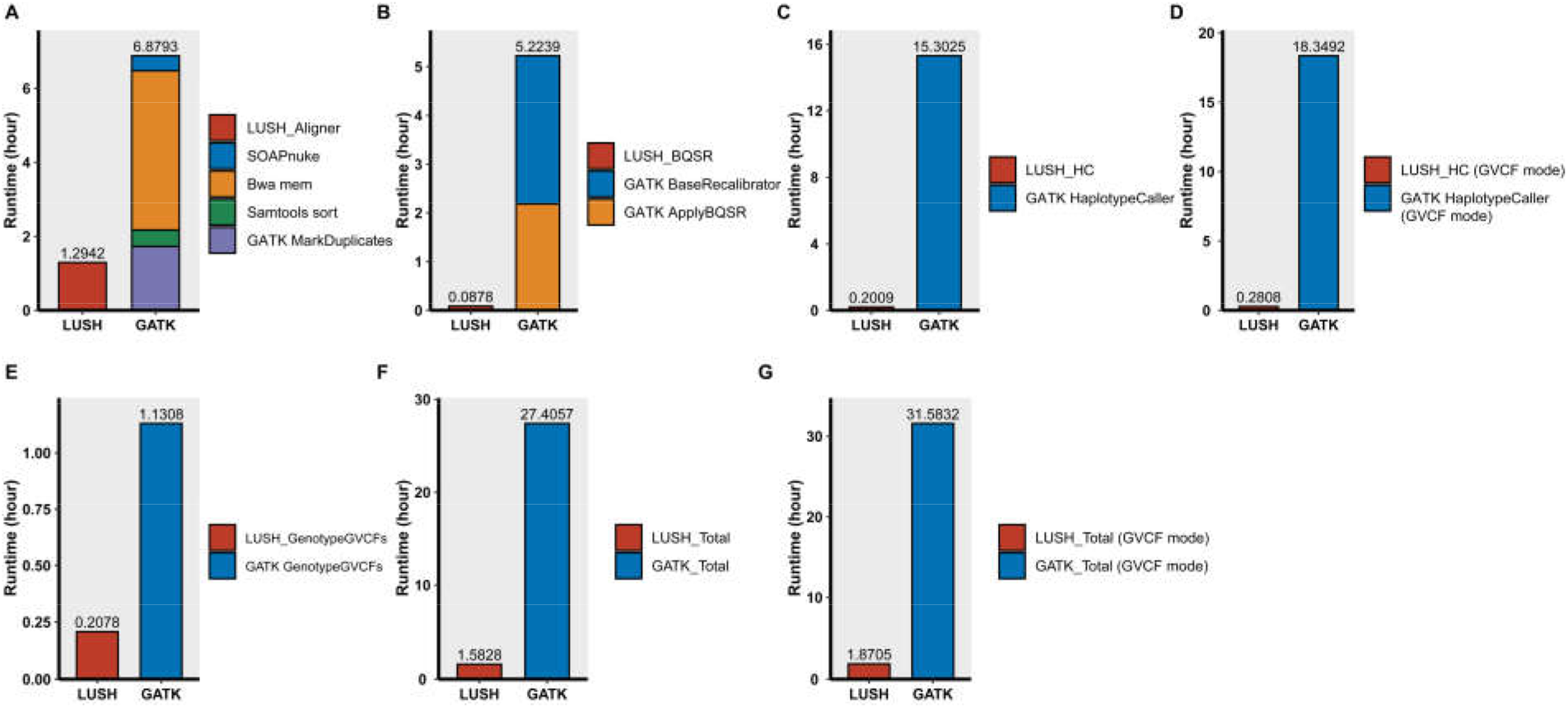
Runtime of LUSH and GATK Variant Calling pipelines on NA12878. **A)** Runtime of each component from FASTQ to BAM. **B)** Runtime of base quality score recalibration. **C)** Runtime of variant calling in non-GVCF mode. **D)** Runtime of variant calling in non-GVCF mode. **E)** Runtime of joint genotyping on one sample. **F)** Total elapsed time in non-GVCF mode. **G)** Total elapsed time in GVCF mode.

The GVCF mode was commonly used in the cohort-wide analysis, which can then be used for joint genotyping of multiple samples in a very efficient way. This enables rapid incremental processing of samples as they roll off the sequencer, as well as scaling to a very large cohort size. Thus, we also implanted GVCF mode in LUSH_HC. In GVCF mode, LUSH_HC used only 0.28 h, while the original GATK pipeline took 18.35 h to process the same dataset (**Figure 3 D**). Regarding the performance of single sample joint genotyping, LUSH_GenotypeGVCFs (0.21h) was 5X faster than GATK-GenotypeGVCFs (1.13h) (**Figure 3 E**).

For the whole pipeline from FASTQ to VCF, the LUSH pipeline greatly reduced the runtime in both non-GVCF and GVCF modes, taking less than 2 hours for 30X WGS data, about 17 times faster than the GATK pipeline (**Figure 3 F, G**). Likewise, the LUSH pipeline had a similar performance on CHM-Synthetic-diploid and two-trios WGS datasets (**Figure S2 A-G, Table S1**).

### Variant Calling accuracy of the LUSH pipeline

We then compared the accuracy of the LUSH pipeline with that of the GATK pipeline. The underlying algorithm of LUSH is roughly the same as that of GATK, so they were expected to produce identical results. We ran each of the three datasets mentioned above using two pipelines. The generated VCFs were compared with their respective truth sets using the haplotype comparison tool hap.py or CHM-evalkit. The comparison was limited to the high-confidence regions of each dataset (see Methodology evaluation). Due to the common use of GVCF mode in cohort studies, we also add GVCF mode to the comparison. As expected, LUSH and GATK demonstrated almost the same precision, recall and F1-score on the NA12878 (**Figure 4 A, B**) and CHM-Synthetic-diploid datasets (**Figure 4 C, D**), both for the SNP and INDEL. Interestingly, the results in non-GVCF mode have higher precision than that in GVCF mode, while the recall is slightly lower. In terms of the F1-Score, the non-GVCF mode performs better (**Figure 4 A, B, C, D**). The findings are also in full agreement with the Two trios WGS data.

**Figure 4.**
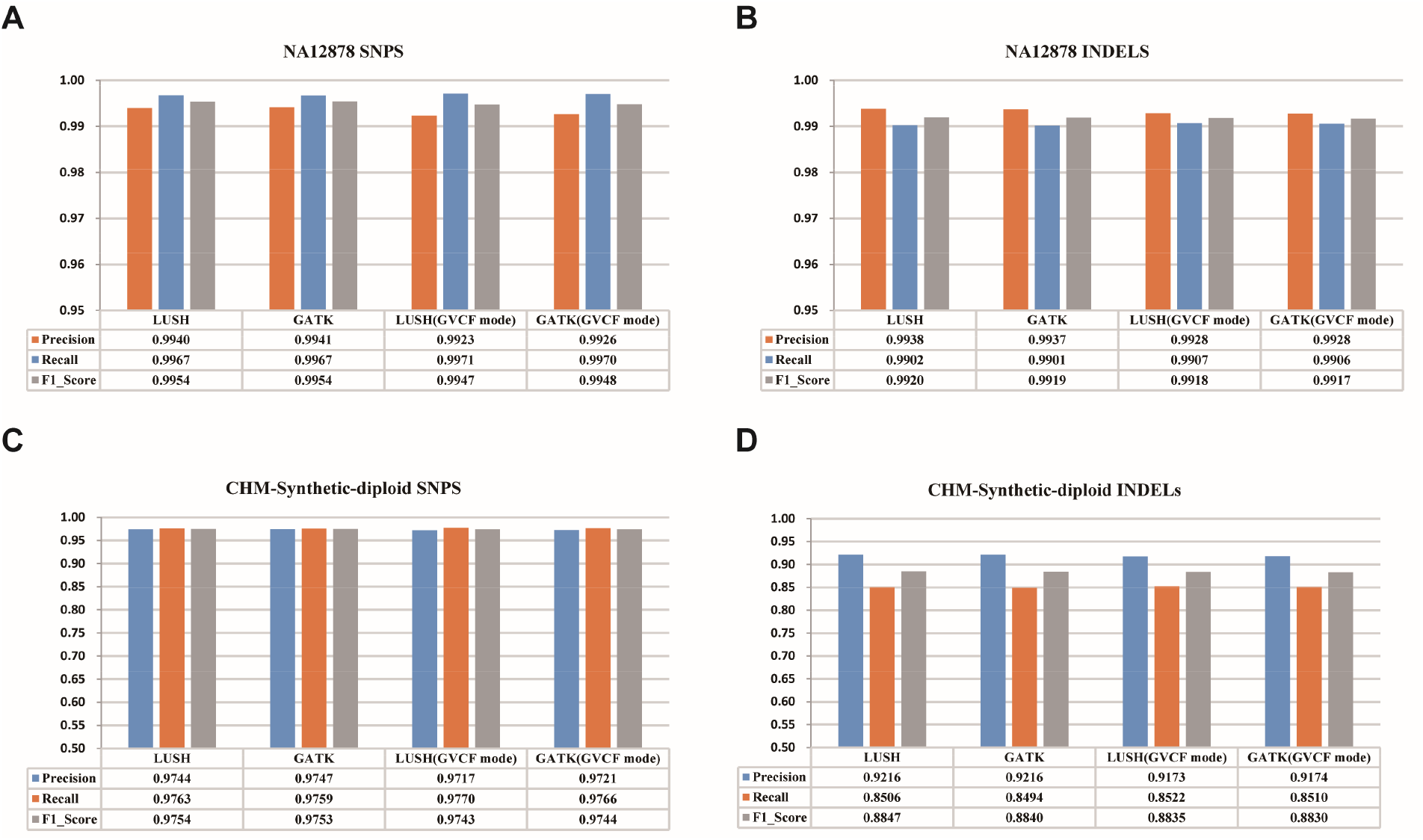
Accuracy of LUSH and GATK Variant Calling pipelines. (A, B) on NA12878. (C, D) on CHM-Synthetic-diploid datasets.

We then analyzed the intersection of the LUSH pipeline and the GATK pipeline. All variants detected by both pipelines were used for the analysis. As expected, approximately 99.11-99.14% of SNPs and 98.92-99.08% of INDELs were co-reported by the two pipelines, both in the non-GVCF mode and GVCF mode, indicating a high consistency of the LUSH and GATK pipelines (**Figure 5 A, B, C, D**). Among these LUSH-only and GATK-only variants, the observed TP rates for SNP were 1.54% and 0.74%, respectively. For INDEL, the TP rates were 5.33% and 2.51%, respectively **(Figure 5 A, B)**. The TP rates in GVCF mode were consistent with these results **(Figure 5 C, D)**. We then annotated genomic regions for LUSH or GATK-specific SNPs and INDELs using SNPEFF. Among these few pipeline-specific variants, LUSH-only and GATK-only variants showed consistent distribution across the genome, and more than 97% of the variants were located in non-functional regions (sum of INTERGENIC, INTRON, DOWNSTREAM, UPSTREAM).

**Figure 5.**
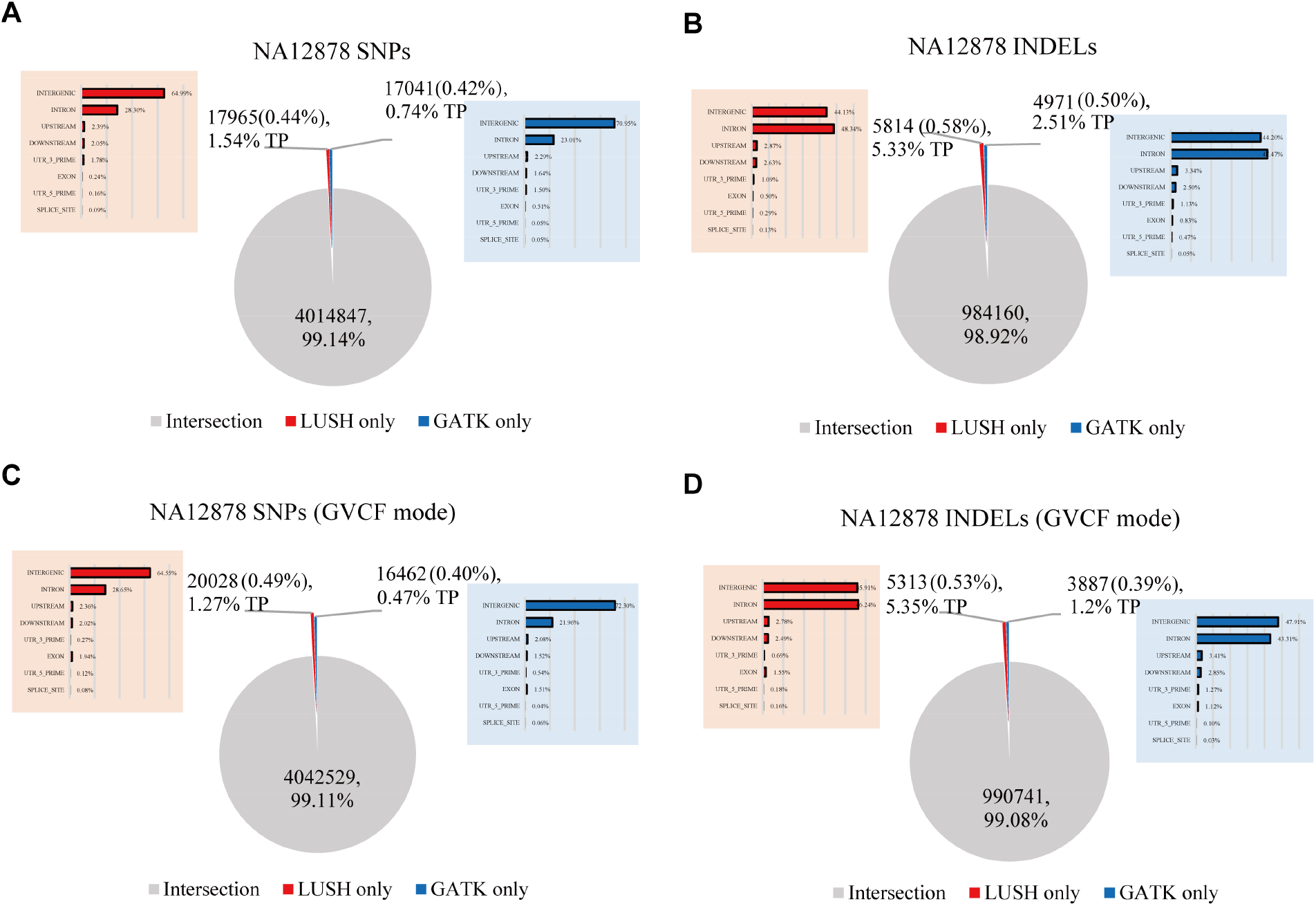
The intersection of variants called by different pipelines on NA12878. (A, B) In non-GVCF mode. (C, D) In GVCF mode.

Moreover, we also explored the intersection of variants in non-GVCF mode and GVCF mode. The results were highly consistent in both modes, with 99.13% (GATK) and 99.25% (LUSH) of co-detected SNPs and 99.11% (GATK) and 99.18% (LUSH) of co-detected INDELs **(Figure S3 A, B, C, D)**. GVCF mode significantly detected more specific SNPs and INDELs.

## Discussion

Genome sequencing has been commonly used for molecular diagnosis, staging, and prognosis, and the massive sequencing data presents a challenge in terms of analysis time. We have developed a LUSH pipeline consisting of LUSH toolkit enabling rapid and precise results. It takes only 1.6 hours to process 30X WGS data from FASTQ to VCF and ~12 minutes from BAM to VCF, with accuracy comparable to GATK, which is extremely critical for acutely ill patients, such as infants in the Pediatric Intensive Care Unit (PICU) and Neonatal Intensive Care Unit (NICU).

We tested the performance of the LUSH pipeline at different thread scales and showed that the LUSH pipeline has remarkable thread scalability. The LUSH component is based on a parallel computing architecture of producer and consumer, which allows it to achieve optimal performance on any machine with a reasonable configuration of parameters. The LUSH pipeline also scales well linearly at different sequencing depths.

LUSH is based on the original WGS “best practices” with a C/C++ implementation that follows the underlying algorithms of the original pipeline. In terms of speed, the optimization of the underlying language, multi-threaded architecture, and algorithmic framework gives the LUSH pipeline an absolute advantage over GATK. Each step in the LUSH pipeline is at least 5 times faster for the same work, and the step LUSH_HC even reaches a speed increase of 76 times. In terms of accuracy, the results of the LUSH pipeline and GATK are equally accurate and highly consistent. The annotation results for specific variants show no meaningful differences in reliability between them. We also demonstrated both high accuracy and hyper speed for the LUSH pipeline on multiple datasets of standards.

## Conclusion

The LUSH workflow composed of the LUSH tools can timely and accurately detect variants from whole-genome sequencing data. This provides important advancements for bedside analysis of acutely ill patients, large-scale analysis of cohort data, and high-throughput variant calling in crop breeding programs. Moreover, LUSH toolkit can be deployed on any CPU-based computing system and have high scalability, which enables it to be quickly applied to various scenarios such as clinical diagnosis and genomic research.

## Declarations

### Ethics approval and consent to participate

All utilized datasets are publicly available. No ethics approval or consent to participate was required for this study.

### Consent for publication

Not applicable.

### Availability of data and materials

All data generated or analyzed during this study are included in this published article and its supplementary information files. Benchmarking scripts and commands for LUSH pipeline are available at https://github.com/Bgi-LUSH/LUSH-DNASeq-pipeline.

### Competing interests

The authors declare that they have no competing interests.

### Funding

Not applicable.

### Authors’ contributions

TW, YZ and HW devised the study, analyzed the data, interpreted the results and drafted the manuscript. TW, JY and QZ conducted the experiments and revised the manuscript. YZ, HW, QZ, TZ, GS, WL, LY, XH and RY implemented the algorithm. CW, ZL and ZL performed data analyses. JW provided critical intellectual comments. XJ and ZH supervised the study and reviewed the manuscript. All authors read and approved the final manuscript.

## Acknowledgements

The authors are grateful to the people in their research group for support and valuable suggestions.

## Supplementary Figures

**Figure S1.**
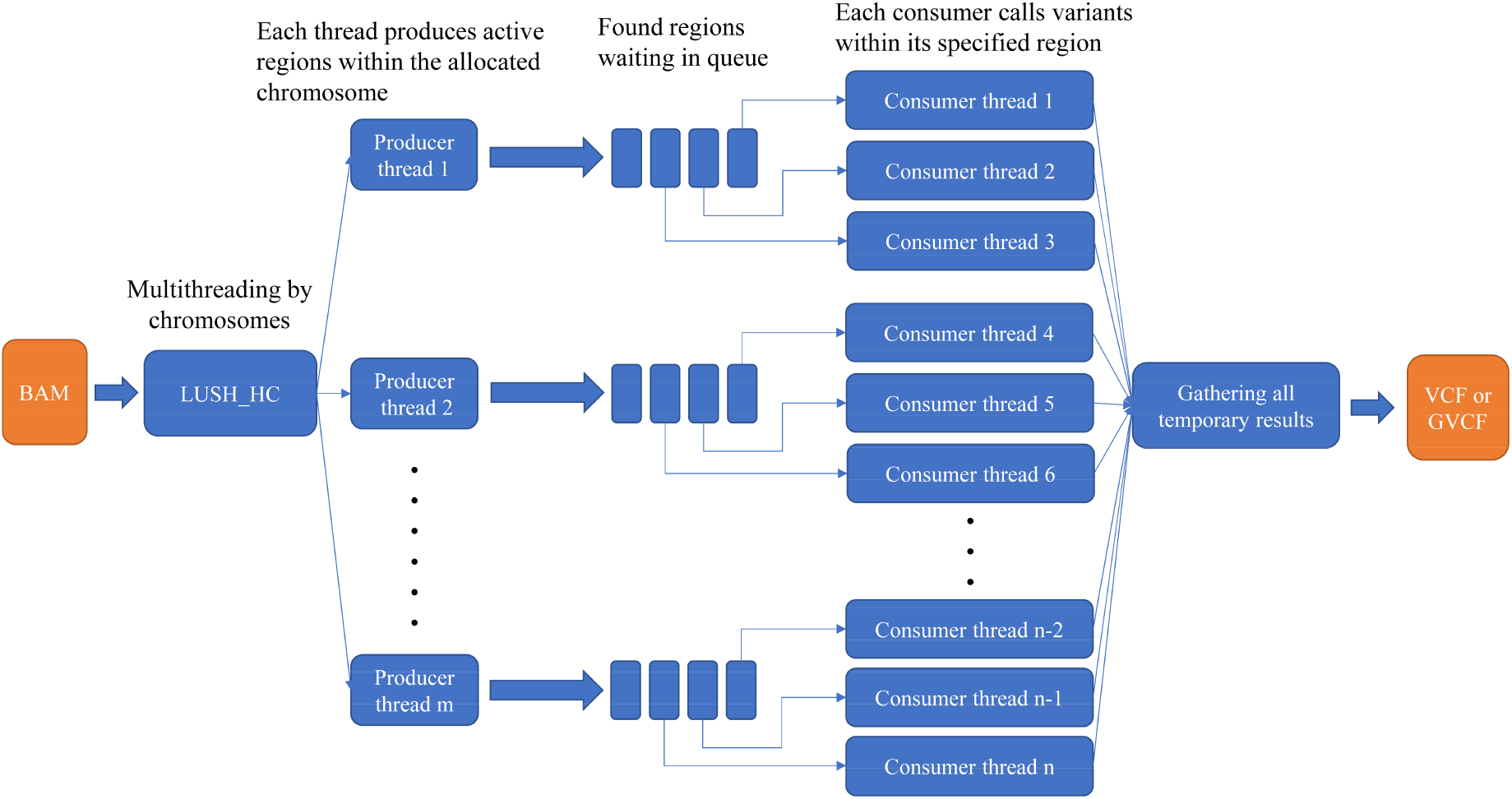
Producer-Consumer pattern in LUSH_HC.

**Figure S2.**
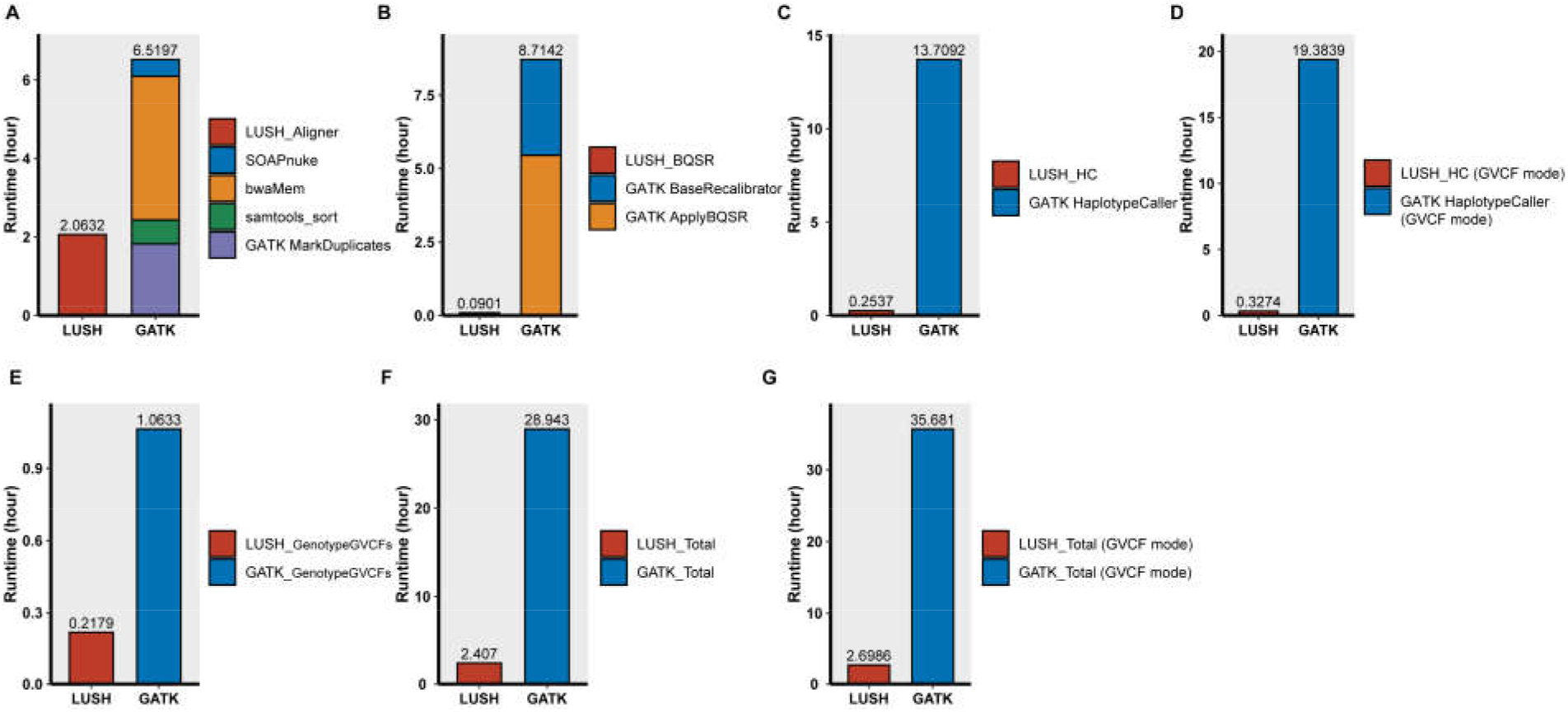
Runtime of LUSH and GATK pipelines on CHM-Synthetic-diploid. **A)** Runtime of each component from FASTQ to BAM. **B)** Runtime of base quality score recalibration. **C)** Runtime of variant calling in non-GVCF mode. **D)** Runtime of variant calling in non-GVCF mode. **E)** Runtime of joint genotyping on one sample. **F)** Total elapsed time in non-GVCF mode. **G)** Total elapsed time in GVCF mode.

**Figure S3.**
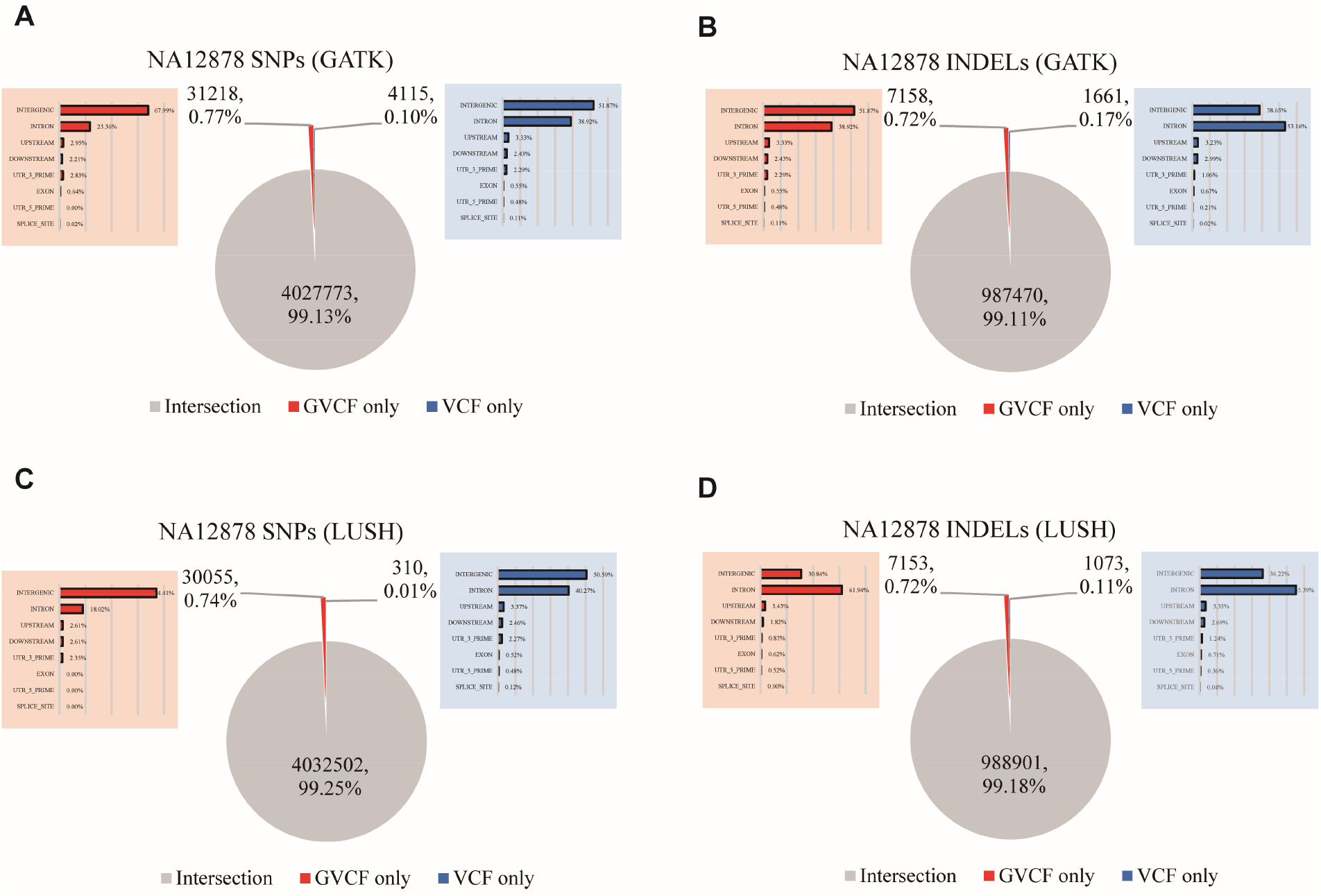
The intersection of variants in non-GVCF mode and GVCF mode. (A, B) For GATK pipeline (C, D) For LUSH pipeline.

## Notes

### Competing Interest Statement

The authors have declared no competing interest.

## References

1. Dewey, F.E., et al., Clinical interpretation and implications of whole-genome sequencing. JAMA, 2014. 311(10): p. 1035–45.

2. Ashley, E.A., et al., Clinical assessment incorporating a personal genome. Lancet, 2010. 375(9725): p. 1525–35.

3. Green, E.D., M.S. Guyer, and I. National Human Genome Research, Charting a course for genomic medicine from base pairs to bedside. Nature, 2011. 470(7333): p. 204–13.

4. Nakagawa, H. and M. Fujita, Whole genome sequencing analysis for cancer genomics and precision medicine. Cancer Sci, 2018. 109(3): p. 513–522.

5. Lupski, J.R., et al., Whole-genome sequencing in a patient with Charcot-Marie-Tooth neuropathy. N Engl J Med, 2010. 362(13): p. 1181–91.

6. Shigemizu, D., et al., Whole-genome sequencing reveals novel ethnicity-specific rare variants associated with Alzheimer’s disease. Mol Psychiatry, 2022. 27(5): p. 2554–2562.

7. Sanford, E., et al., Clinical utility of ultra-rapid whole-genome sequencing in an infant with atypical presentation of WT1-associated nephrotic syndrome type 4. Cold Spring Harb Mol Case Stud, 2020. 6(4).

8. Xing, R., et al., Whole-genome sequencing reveals novel tandem-duplication hotspots and a prognostic mutational signature in gastric cancer. Nat Commun, 2019. 10(1): p. 2037.

9. Bick, D., et al., Case for genome sequencing in infants and children with rare, undiagnosed or genetic diseases. J Med Genet, 2019. 56(12): p. 783–791.

10. Saunders, C.J., et al., Rapid whole-genome sequencing for genetic disease diagnosis in neonatal intensive care units. Sci Transl Med, 2012. 4(154): p. 154ra135.

11. Willig, L.K., et al., Whole-genome sequencing for identification of Mendelian disorders in critically ill infants: a retrospective analysis of diagnostic and clinical findings. Lancet Respir Med, 2015. 3(5): p. 377–87.

12. McKenna, A., et al., The Genome Analysis Toolkit: a MapReduce framework for analyzing next-generation DNA sequencing data. Genome Res, 2010. 20(9): p. 1297–303.

13. Heldenbrand, J.R., et al., Performance benchmarking of GATK3. 8 and GATK4. BioRxiv, 2018: p. 348565.

14. Pluss, M., et al., Need for speed in accurate whole-genome data analysis: GENALICE MAP challenges BWA/GATK more than PEMapper/PECaller and Isaac. Proc Natl Acad Sci U S A, 2017. 114(40): p. E8320–E8322.

15. Raczy, C., et al., Isaac: ultra-fast whole-genome secondary analysis on Illumina sequencing platforms. Bioinformatics, 2013. 29(16): p. 2041–3.

16. Kendig, K.I., et al., Sentieon DNASeq Variant Calling Workflow Demonstrates Strong Computational Performance and Accuracy. Front Genet, 2019. 10: p. 736.

17. Miller, N.A., et al., A 26-hour system of highly sensitive whole genome sequencing for emergency management of genetic diseases. Genome Med, 2015. 7: p. 100.

18. Chen, Y., et al., SOAPnuke: a MapReduce acceleration-supported software for integrated quality control and preprocessing of high-throughput sequencing data. Gigascience, 2018. 7(1): p. 1–6.

19. Li, H. and R. Durbin, Fast and accurate short read alignment with Burrows-Wheeler transform. Bioinformatics, 2009. 25(14): p. 1754–60.

20. Li, H., et al., A synthetic-diploid benchmark for accurate variant-calling evaluation. Nat Methods, 2018. 15(8): p. 595–597.

21. Cingolani, P., et al., A program for annotating and predicting the effects of single nucleotide polymorphisms, SnpEff: SNPs in the genome of Drosophila melanogaster strain w1118; iso-2; iso-3. Fly (Austin), 2012. 6(2): p. 80–92.

22. Danecek, P., et al., Twelve years of SAMtools and BCFtools. Gigascience, 2021. 10(2).

